# A crocodylian-style cloaca in a non-avialan dinosaur

**DOI:** 10.1101/2020.10.11.335398

**Authors:** Phil R. Bell, Michael Pittman, Thomas G. Kaye, Christophe Hendrickx

## Abstract

Our knowledge of the reproductive biology of dinosaurs covers a range of aspects, from brooding behaviour to nesting style and the timing of sexual maturity. Yet, the basic anatomy and function of the cloaca in non-avialan dinosaurs remains unknown. Here, we describe the outer morphology of the only known non-avialan dinosaur cloaca, preserved in an exceptional specimen of the early-diverging ceratopsian dinosaur *Psittacosaurus*. We clarify the position of the cloaca with respect to the ischia and caudal vertebrae and document the scales immediately adjacent to the abdomen and tail. We find that the cloaca is from a near-sexually mature subadult individual and is most similar to the cloaca of crocodylians, to the exclusion of lepidosaurians and birds. However, the sex of SMF R 4970 could not be determined as the cloaca and the rest of the specimen does not yield any sexually dimorphic information. This study highlights the ongoing role of exceptional specimens in providing rare soft tissues that help to bridge longstanding gaps in our knowledge of the basic biology of dinosaurs and other extinct reptiles.

## Introduction

The reproductive biology of extinct non-avialan dinosaurs is rarely interpreted from the fossil record. To date, exceptionally well-preserved remains and the extant phylogenetic bracket (EPB; [1]) have clarified details including their brooding behaviour, nesting style and timing of sexual maturity [2-7]. However, the anatomy and function of the cloaca has continued to remain elusive. In archosaurian and lepidosaurian reptiles, the cloaca forms the common opening of the digestive and urogenital tract and consists of a series of chambers—the coprodeum, urodeum, and proctodeum—separated by muscular sphincters, and which terminates in the vent [8-10]. As a result, it serves as the passage for digestive and urinary wastes, transmission of the male copulatory organ, and the passage of eggs or live young in addition to a variety of less obvious functions such as salt and temperature regulation [11]. Nearly 70 years ago, Alfred Romer predicted that the cloaca in extinct archosaurs was positioned within the proximal tail region where haemal arches are absent [1]. Here, we provide confirmation of Romer’s original prediction based on an exquisitely-preserved specimen of an early-diverging ceratopsian dinosaur, *Psittacosaurus* sp., discovered from the Early Cretaceous Jehol Group of northeastern China [12]. This specimen, known for its exquisite coloured epidermal structures and elongate caudal monofilaments [12-14], preserves for the first time the outer morphology of the cloaca in a non-avialan dinosaur.

## Results

### Age of the individual

The right femur of Senckenberg Museum Frankfurt (SMF) R 4970 is ∼140mm long. This is similar to the femoral lengths of *P. lujiatunensis* Institute of Vertebrate Paleontology and Paleoanthropology (IVPP) V12617 (138mm) and V18344 (145mm), Liaoning Paleontological Museum (LPM) R00128 (135mm) and R00138 (144mm) and Peking University Vertebrate Paleontology (PKUVP) V1053 (149mm) and V1056 (135mm) which belong to ∼6-7 year old subadults (see Table 1 and Fig. 5 of Erickson *et al*. [15] and Supplementary Table S2 of Zhao *et al*. [16]). This age is just shy of sexual maturity and at the beginning of the exponential growth phase (see Table 1 and Fig. 5 of Erickson *et al*. [15] and Supplementary Table S2 of Zhao *et al*. [16]). The femoral length of SMF R 4970 is therefore the closest match to a nearly sexually mature subadult (see Table 1 and Fig. 5 of Erickson *et al*. [15] and Supplementary Table S2 of Zhao *et al*. [16]). Thus, the configuration and morphology of the cloaca in SMF R 4970 and its surrounding osteological correlates had not yet reached its most developed form or full sexual maturity.

### The cloaca and its surrounding scales and bones

Immediately posterior to the ischiadic symphysis is the fleshy aperture (or vent) of the cloaca (Figs. 1, 2). Since SMF R 4970 is preserved lying on its back, both the left and right sides of the cloaca are visible, although the right side is better exposed (Fig. 1). The vent itself is longitudinally oriented (∼2 cm long) and surrounded on either side by darkly pigmented tissue. This appears to lie flush with the ventral parts of the abdomen and tail (i.e. it does not form a cloacal protuberance). The darker cloaca is clearly differentiated, from the lighter-coloured surrounding integument (Fig. 2).

**Figure 1.**
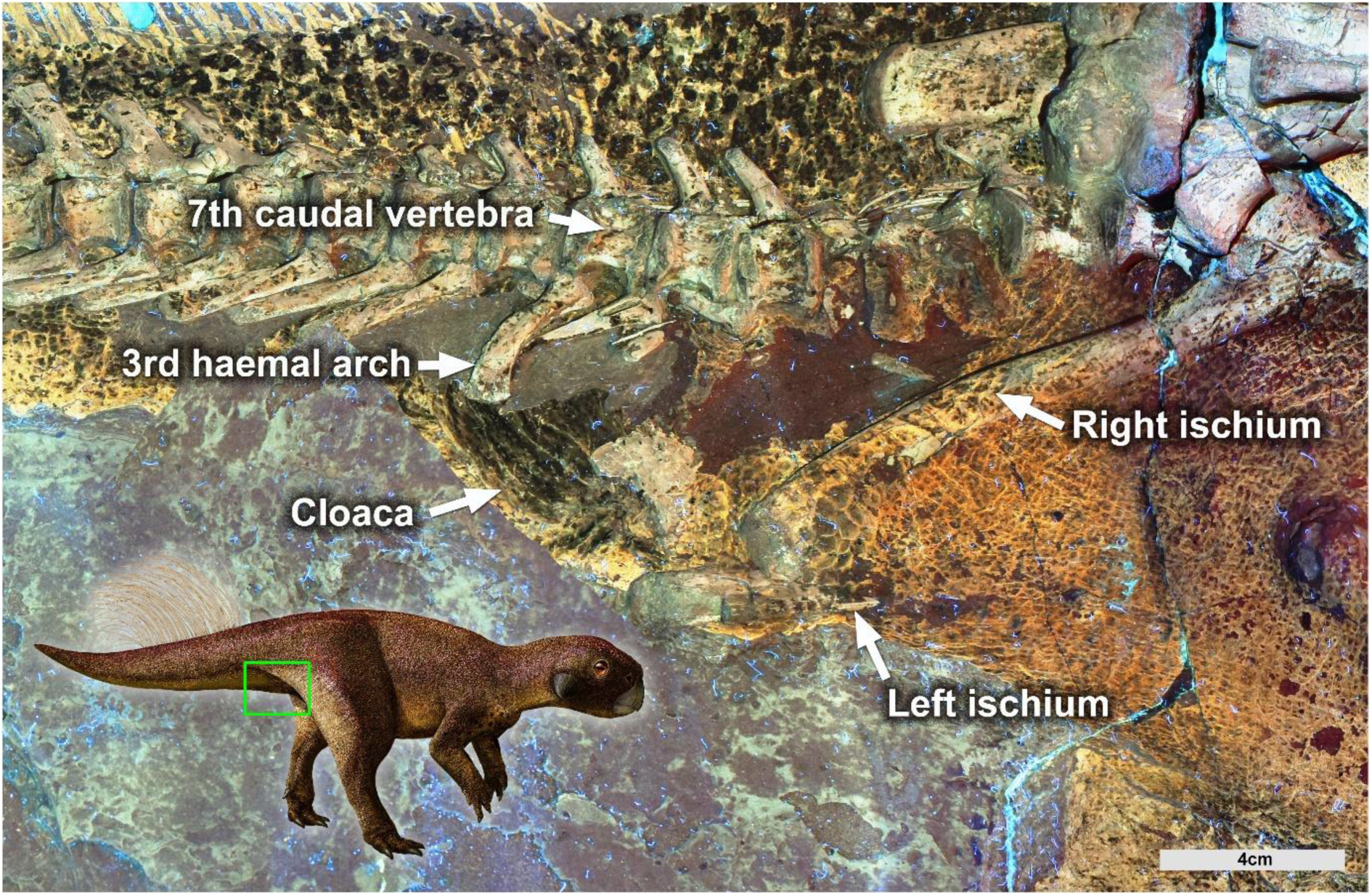
Cloacal region of the ceratopsian dinosaur *Psittacosaurus* under Laser-Stimulated Fluorescence (LSF). The cloaca soft tissue is the blackish mottled ovoid area preserved in the region between the ischia and the third haemal arch in specimen SMF R 4970 from the Early Cretaceous Jehol Group of northeastern China. Scale is 4cm. Life reconstruction with green box showing the position of the LSF image. Image Credit: Julius T. Csotonyi (image used with permission).

**Figure 2.**
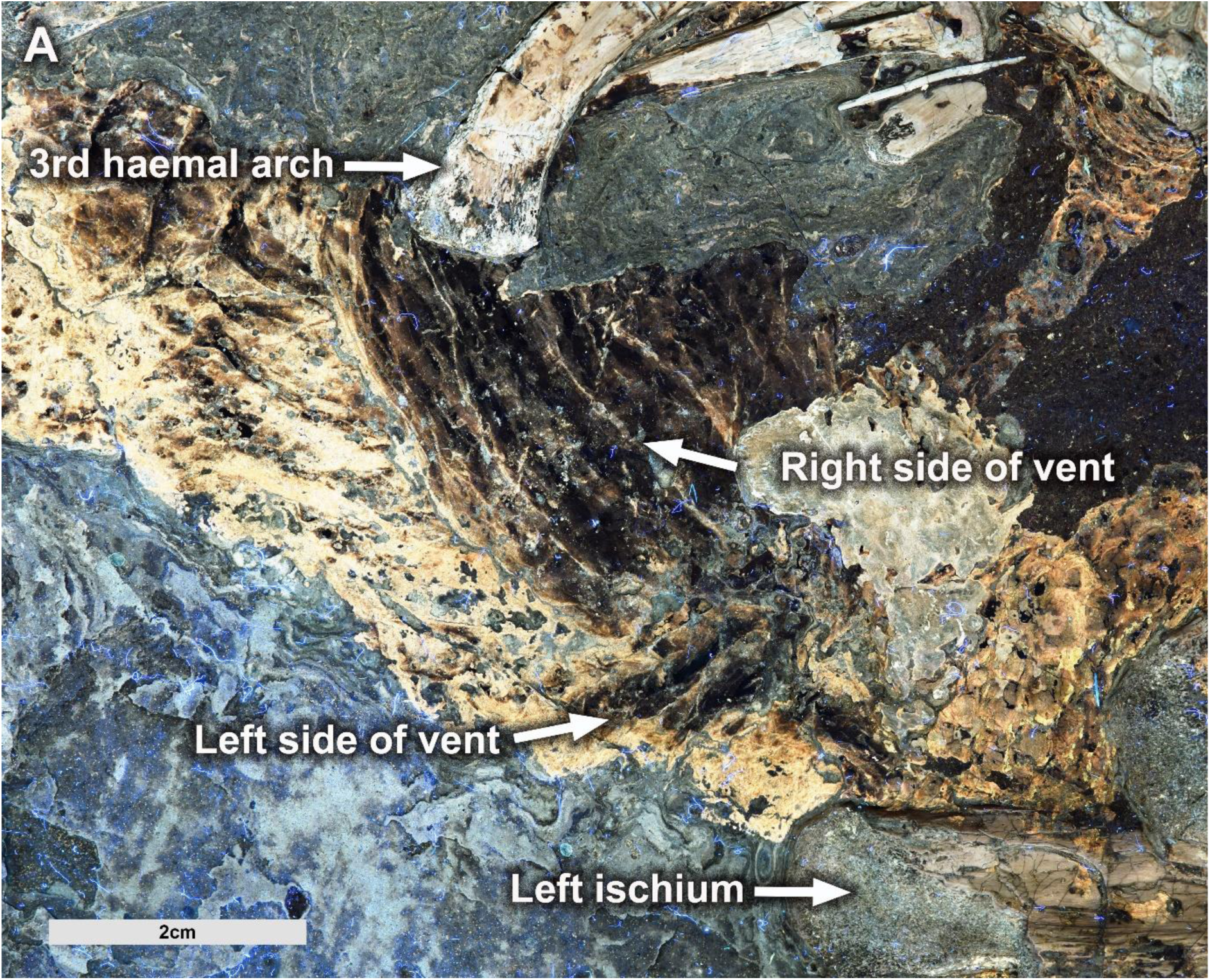
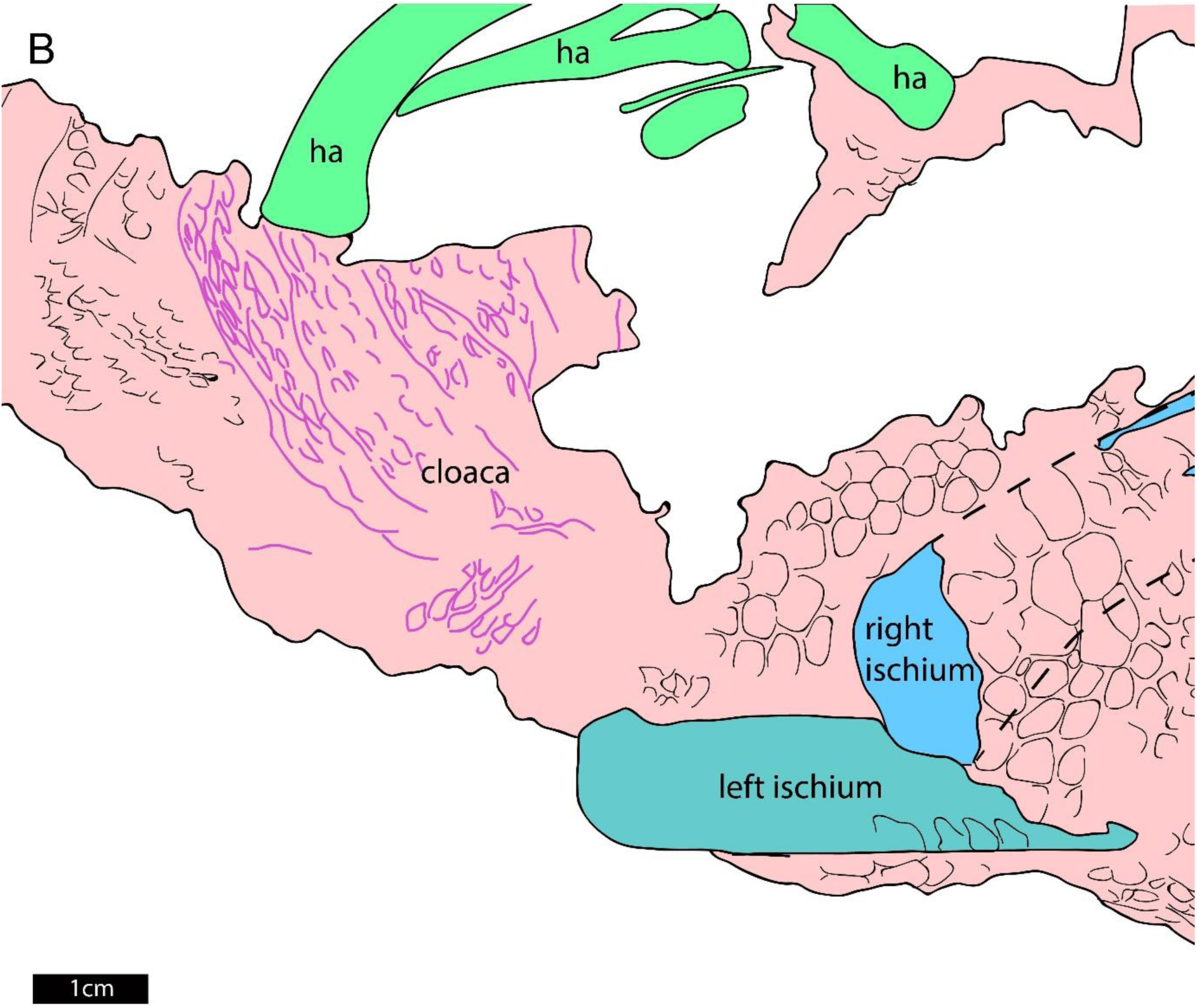
Close up of the cloaca of *Psittacosaurus* sp. SMF R 4970. **A**. The right side of the longitudinally presented vent is better exposed. The outer morphology of the cloaca is darkly pigmented compared to adjacent areas of the ventrum. Scale bar is 2cm. **B**. Interpretative drawing of the cloacal region showing the integument in red shading and the details of the cloaca in pink line colour. Haemal arches (ha) marked in lime green and left and right ischia coloured in dark green and blue, respectively. Scale bar is 1 cm.

The darkly-pigmented tissue surrounding the vent is wrinkled, the creases of which are parallel and radiate from the vent in a posterolateral direction (∼3 cm long) towards the ventral tip of the third haemal arch (Fig. 2). Unlike the other haemal arches, the third haemal arch, which articulates with the posteroventral corner of the 7^th^ caudal vertebra, is subvertical and strongly bowed (anteriorly concave) (Fig. 1). In contrast, the remaining haemal arches are all straight and strongly posteroventrally oriented (Fig. 1). The first and second haemal arches are also peculiar in being roughly half the length of the third arch, although the first arch appears to be broken into two adjacent lengths (Fig. 1). The scales covering the darkly pigmented tissue surrounding the vent are also distinct from those on the abdomen and tail: scales surrounding the vent are relatively large, roughly lenticular (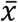 length = 3.4 mm), the long axes of which are oriented parallel to the surrounding wrinkles (Figs. 1, 2). Further along the tail, the ventral integument consists of distinct vertical bands (arranged transversely in life) of typically rounded-quadrangular scales, which range from 1.7 to 3.3 mm in height (Fig. 1). The same transverse banding of quadrangular scales also occurs on the abdomen, anterior to the ischiadic symphysis (Fig. 1).

## Discussion

### Cloaca anatomy and biological implications

The *Psittacosaurus* specimen SMF R 4970 is unique among non-avialan dinosaurs in preserving the outer morphology of the cloaca, including the vent. SMF R 4970 demonstrates unequivocally that the genitals were housed internally in *Psittacosaurus* and likely in all other non-avialan dinosaurs, as in all known animals with a cloaca. For comparison with the cloaca of SMF R 4970, a representative sample of snakes, lizards, birds, and crocodylians were observed in the collections of the University of New England’s Natural History Museum. The internal anatomy of the cloaca in lepidosaurians, crocodylians, and birds differs substantially and is accompanied by modifications to the external appearance of the vent. The vent in living sauropsids can be divided into three main types [17]: transversely opening (snakes and lizards), longitudinally opening (crocodylians), or round/square (birds). The integumentary covering across these three types differs accordingly. In snakes, the transverse vent is covered by one or two cloacal scales that are modifications of the broad ventral scales present elsewhere on the underbelly. The condition in lizards is highly variable but the vent is always transverse and accompanied by a variable number of cloacal scales that may or may not differ from the surrounding scales. Among birds, the area immediately surrounding the cloaca is naked, bearing neither scales nor feathers. In crocodylians, the longitudinal vent is surrounded by elliptical-to-polygonal scales that radiate and increase in size from the vent itself. This rosette arrangement of cloacal scales is distinct from the transverse rows of comparatively large quadrangular scales that extend along the ventral surfaces of the abdomen and tail. Thus, the gross morphology of the vent in *Psittacosaurus*, which combines a longitudinally opening vent with a rosette pattern of cloacal scales and transverse rows of quadrangular ventral scales, most closely matches that of crocodylians.

The position of the vent is also distinctive in *Psittacosaurus*. Romer [1] remarked that the cloaca opens immediately posterior to the distal ischia in all reptiles; however, it may be supported by an ossified hypoischium that projects posteriorly from the ischial symphysis in some lizards (see [18] and references therein). In many lizards, the vent is typically positioned just posterior to the joint between the first and second caudal vertebrae and is never further than the third caudal vertebra [18]. Birds on the other hand have highly modified tails that terminate in a pygostyle (e.g. [19]) and therefore do not provide a suitable analogue for comparisons. Matching Romer’s [1] prediction, the vent, in *Psittacosaurus*, opens just posterior to the distal ischia and in line with the joint between the fifth and sixth caudal vertebrae.

The crocodilian-like vent of *Psittacosaurus* implies that, unlike lizards and later-diverging birds, *Psittacosaurus* probably had a muscular, unpaired, and ventrally-positioned copulatory organ (e.g. [20]) and a ureter that was decoupled from the copulatory organ [9]. Some birds possess a single copulatory organ, but the majority of species lack a phallus entirely [21]. Like crocodylians, birds still use internal fertilisation, which is the presumed method in *Psittacosaurus*. Crocodylians and some early-diverging crown birds (e.g., *Rhea*, tinamous) have a ureter that opens into the coprodeum, whereas in later-diverging crown birds, the ureter empties into the urodeum [9]. The presumably paired oviducts in *Psittacosaurus* [22] would have opened into the urodeum as well.

Despite the remarkable preservation of SMF R 4970, it was not possible to determine its sex. Although histological examination was not available, SMF R 4970 was not gravid, and therefore unlikely to have been producing medullary bone [23]. There is no evidence of a recognisable sexually dimorphic cloacal structure in SMF R 4970. A cloacal protuberance temporarily develops in some birds during the breeding period, but its utility as a dimorphic structure is undermined by its presence in both males and females in some species [24]. In crocodylians, sex determination is entirely dependent on the inspection of the genitalia and has no relationship to the external morphology of the cloaca/vent [25, 26]. Internally, a reduced first (and second) haemal arch—as in SMF R 4970—has been suggested to identify female dinosaurs based on the alleged condition in modern crocodylians [1, 27]. However, this hypothesis has no statistical support [28]. The enlarged third haemal arch was likely the origin for muscles responsible for closing the cloaca (m. transversus medianus [17]). Anterior haemal arches are commonly abbreviated in dinosaurs and seem to be an ancestral feature based on rare early dinosaur fossils preserving this portion of the tail (e.g. *Chilesaurus, Eoraptor*, and *Tawa*). Their presence in modern crocodylians suggests they might even be ancestral to archosaurs pending further fossil tail discoveries. Thus, the only known record of a non-avialan dinosaur cloaca has a crocodylian-style and does not show external sexually dimorphic structures that can determine the sex of SMF R 4970.

## Methods

SMF R 4970 was photographed using Laser-Stimulated Fluorescence (LSF) performed using an updated version of the methodology proposed by Kaye *et al*. [29] and refined in Wang *et al*. [30]. Only a brief description of the method is provided here. A 405 nm violet near-UV laser diode was used to fluoresce the specimen following standard laser safety protocol. Long exposure images were taken with a Nikon D810 DSLR camera fitted with a 425 nm long-pass blocking filter and controlled from a laptop using *digiCamControl*. Image post-processing (equalization, saturation and colour balance) was performed uniformly across the entire field of view in *Photoshop CS6*. As soft tissue details in SMF R 4970 are best visible under LSF, the observations made using this technique and the resulting digital images formed the basis for our descriptions. All measurements were taken using digital images uploaded and calibrated in *ImageJ v1*.*52q* or *Photoshop CS6*. Comparative anatomy involved a representative sample of snakes, lizards, birds and crocodylians, which were observed in the collections of the University of New England’s Natural History Museum (Armidale, Australia). Computed tomography (CT) scans of a subadult male *Crocodylus porosus* scanned and acquired by Klinkhamer *et al*. [31] were visualised using *Dragonfly 2020*.1 (www.theobjects.com).

## Acknowledgements

We thank Rainer Brocke, Olaf Vogel, and Gerald Mayr (SMF) for specimen access, J. Holley (UNE) for access to comparative collections, and T. Frauenfelder (UNE) for assistance with *Dragonfly 2020*. P.R.B. was supported by an Australian Research Council Discover Early Career Research Award (project ID: DE170101325). M.P. and T.G.K. are supported by a RAE Improvement Fund of the Faculty of Science, The University of Hong Kong (HKU) and HKU MOOC course Dinosaur Ecosystems. C.H. is supported by the Consejo Nacional de Investigaciones Científicas y Técnicas (CONICET), Argentina (Beca Pos-doctoral Legajo 181417).

